# Prediction and interpretation of deleterious coding variants in terms of protein structural stability

**DOI:** 10.1101/210120

**Authors:** F. Ancien, F. Pucci, M. Godfroid, M. Rooman

## Abstract

The classification of human genetic variants into deleterious and neutral is a challenging issue, whose complexity is rooted in the large variety of biophysical mechanisms that can be responsible for disease conditions. For non-synonymous mutations in structured proteins, one of these is the protein stability change, which can lead to functionality loss. We developed a stability-driven knowledge-based classifier that uses protein structure, artificial neural networks and solvent accessibility-dependent combinations of statistical potentials to predict whether destabilizing or stabilizing mutations are disease-causing. Our predictor yields a balanced accuracy of 71% in cross validation. As expected, it has a very high positive predictive value of 89%: it predicts with high accuracy the subset of mutations that are deleterious because of stability issues, but is by construction unable of classifying variants that are deleterious for other reasons. Its combination with an evolutionary-based predictor increases the balanced accuracy up to 75%, and allowed predicting more than 1/4 of the deleterious variants with 95% positive predictive value. Our method, called SNPMuSiC, can be used with both experimental and structural models and compares favorably with other prediction tools on several independent test sets. It constitutes a step towards interpreting variant effects at the molecular scale.

## Introduction

Despite the large amounts of genomic data collected in the last decade and the multiple efforts to elucidate their links with phenotypic traits, an accurate and interpretative classification of the effects of genetic variants on various disorders remains a difficult goal to achieve. The high complexity of the problem is mainly due to the heterogeneity of the molecular mechanisms underlying the diseases, most of which have still to be deeper understood to be useful at the clinical level. For example, a change in a single DNA base pair that occurs in a coding region can be synonymous, i.e. change the codon but not the amino acid, and perturb the gene expression level or the efficiency of the translational mechanism. Non-synonymous mutations that change both the codon and the associated amino acid can modify the biophysical properties of the encoded protein, such as its thermodynamic stability or solubility, or its functional properties by affecting the active site or the binding affinity for ligands and proteins partners. Moreover, the impact of a given variant crucially depends on its context within the genome: its deleteriousness can depend on the presence of other variants in the same or in other genes. Its effect also varies according to the cellular context. Indeed, some variants occur in proteins that play an essential role for the cell or the organism, which cannot be performed by any other protein. In such case, even a slightly destabilizing variant can be strongly deleterious. In contrast, some proteins perform functions that can very well be performed by other proteins, so that inactivating mutations do not cause diseases.

Among all the possible types of variants, the non-synonymous single nucleotide variants in coding genes (SNV) play an important role since they constitute more than half of the mutations known to be associated in human inherited disorders.^1^ They are directly related to a wide range of pathological conditions among which Parkinson’s and Alzheimer’s diseases, and are involved in complex diseases such as cancers, in which the accumulation of different types of genetic variants determine the tumor initiation and progression.^2^

It is thus of primary interest, especially in the context of personalized medicine, to have computational methods to classify SNVs and identify those that are disease causing. Classification methods developed in the literature usually integrate different kinds of biological data in view of gathering the complexity of the phenomenon, and require only the amino acid sequence as input. Some of them, such as Provean,^3,4^ Sift^11^ or the Evolutionary Action method,^5,6^ use as sole ingredient the evolutionary conservation in homologous proteins at the mutated position and in the neighborhood along the sequence. Though this information is represented by a single score, it corresponds to a mixture of several molecular effects, among which protein stability, solubility, function, and interactions. Understanding why a residue is conserved is impossible on the basis of the evolutionary data only. Other methods add further layers of information on top of this data, by considering for example structural alterations in the biophysical characteristics of proteins, which are usually predicted from the sequence and sometimes derived from the structure when available; these methods include Mutation Tasser,^7^ Mutation Assessor,^8^ CADD,^9^ Polyphen-2,^10^ DUET,^13^ SuSPect,^14^ and VIPUR.^12^ The addition of contextual information about the protein-protein interaction networks, the protein essentiality index, and the pathway in which the protein is involved appear to significantly improve the performances, as shown in DEOGEN.^16,17^ These various computational tools use machine learning techniques to predict from the considered features the effect of missense mutations.

Unfortunately, the accuracy of the prediction methods remain limited, with often a quite high false-positive rate with the detrimental consequence that many of the predicted deleterious variants observed in clinical whole-exome sequencing turn actually out to be neutral.^18^ Frequently moreover, the computational methods only focus on reaching the highest variant classification accuracy on a given dataset rather than predicting and understanding the modifications that occur at the molecular scale and are responsible for the altered phenotype; yet this information is a prerequisite for the rational design of drugs or treatments. As a matter of fact, although protein structure can help getting insight into in the molecular impact of mutations, it is rarely (fully) exploited by variant classification methods, except in a few cases.^12,13^,^15,19^,^20^

The present analysis aims at deepening the understanding of the relation between protein structural stability and variant deleteriousness. To evaluate the stability of a given protein structure and its change upon single-site mutations, we used knowledge-based statistical mean-force potentials derived from a dataset of three-dimensional (3D) protein structures. Combinations of these potentials, performed with the help of different artificial and probabilistic neural network architectures that include the solvent accessibility of the mutated residue to modulate the importance of the energetic terms, were used to classify SNVs into neutral and deleterious. Note that the primary goal of this analysis is not to reach the highest prediction accuracy but rather to get insight into the functional and stability characteristics of protein variants and their relation with the phenotypic traits. Our objective also involves predicting with high accuracy the subset of mutations that are deleterious due to stability problems.

## Methods

### Non-synonymous SNV datasets for training and testing

The training dataset was built by combining the annotated, non-synonymous SNV data from three different databases: DbSNP,^21^ SwissVar^22^ and HumSaVar.^23^ In a first stage, we combined all SNVs while avoiding repeats. Note that variants occurring in more than one database with different neutral/deleterious annotation were discarded. The SNVs were characterized in these databases by the protein’s UniProt code^23^ and the variant’s residue number, without any reference to a possible 3D structure. We had thus, in a second stage, to identify the subset of variants introduced in proteins that have an experimental structure, using the SIFTS webserver.^24^ We only considered the subset of mutations introduced in:

- globular proteins or proteins located in the cytoplasmic or extracellular domain of membrane proteins;
- proteins with an X-ray structure available in the Protein DataBank^25^ (PDB) of resolution of 2.5 Å at most.

Some of the PDB protein structures have not exactly the same sequence as the corresponding entry in the UniProt database, but contain one or several mutations. We only retained the subset of SNVs of residues that are far from all these additional mutations by an inter-C*α* distance of 10 Å at least, to avoid direct interactions between them. The final dataset is referred to as **S** and is reported in Table S1 of Supplementary Material. It contains 5,302 variants inserted in 1,016 different proteins, 1,301 of which are annotated as polymorphic and 4,001 as deleterious.

To test our predictors and compare their performances with other commonly used classification tools, we applied them to four test sets: **S**_**H**_ (variants in Human proteins), **S**_**NH**_ (in Non-Human proteins), **S**_**CV**_ (from the ClinVar dataset^26^) and **S**_**SSC**_ (from the Simon Simplex Collection^27^), which were obtained from^12^ and are described in Supplementary Material (Tables S2-S5).

### Statistical Potentials and other structural features

We used the statistical potential formalism to evaluate the stability of a protein and its change caused by non-synonymous SNVs, and to understand the link with their polymorphic or pathogenic effect. These potentials are knowledge-driven mean force potentials that are extracted from a dataset of well-resolved X-ray protein structures.^28–32^ The energetic contribution *ΔW* associated to the sequence-structure association (*s, c*), where *s* and *c* are sequence and structure elements respectively, is obtained from the inverse Boltzmann law as:

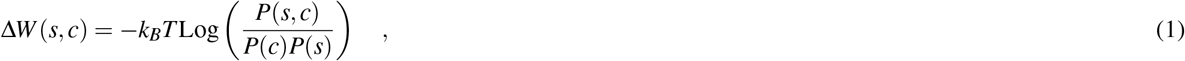

where k_B_ is the Boltzmann constant, *T* the absolute temperature, and *P*(*c, s*), *P*(*c*) and *P*(*s*) are the probabilities of observing (*c, s*), (*c*) or (*s*) elements. These probabilities are approximated in terms of the number of occurrences of these elements in a reference dataset of protein structures. The sequence elements *s* are amino acid types and the structural motifs *c* can be interresidue distances, torsion angle domains or solvent accessibilities. Higher order potentials, in which different combinations of sequence and structure elements are considered, have also been utilized in this investigation. For example, a 3-terms potential describing the association between one structure and two sequence elements is defined as:

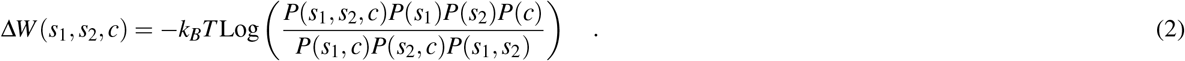

For sake of simplicity, such potentials are referred to as *ΔW*_*ssc*_ in what follows. Further generalizations and technical details can be found in.^32,33^

We used here 13 statistical potentials, listed in Table S6, which can be grouped in three classes according to the type structural descriptor *c*: (1) distance potentials describing tertiary interactions, in which *c* is the distance *d* between the average side chain geometric centers of two amino acids; (2) solvent accessibility potentials, in which *c* is the solvent accessibility *a* of a given residue; (3) torsion potentials describing local interactions along the sequence, in which *c* is the main chain torsion angle domain *t* of a residue. Several combinations of these basic potentials were also used. Besides these potentials, we also considered two biophysical characteristics: the solvent accessibility *A* of the mutated residues and the difference in volume Δ*V* between the wild type and the mutated residues.^33^

These potentials and structural features were used to estimate, for each mutation of our SNV dataset **S**, the corresponding change in volume Δ*V* and in folding free energy ΔΔ*W*_*i*_, with *i* = 1, …13. In a first step, the distributions of ΔΔ*W*_*i*_, *A* and Δ*V* were compared between deleterious and neutral variants from **S**, in view of getting insight into the relation between the structure and stability signals and the pathogenicity of the variants. In a second stage, all the potentials and features were combined to set up a predictor of variant deleteriousness.

### Utilizing neural network architectures for variant classification

#### Pathogenicity prediction from thermodynamic and thermal stability changes

The first approach for predicting on a structural basis which variants are disease-causing and which are not consists in using our PoPMuSiC^33^ and HoTMuSiC^35^ algorithms, which predict changes in thermodynamic and thermal stabilities upon single-site mutations from protein 3D structures. More precisely, PoPMuSiC uses a combination of the 13 statistical potentials and the two structural features listed in Table S6 to estimate the values of the folding free energy changes upon mutation:

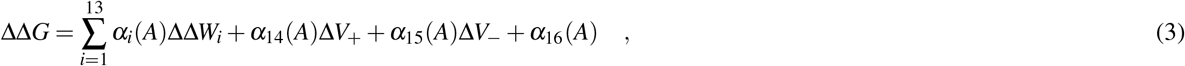

where *ΔV*_+_ and Δ*V*_−_ are two volume terms defined as: *ΔV*_±_ = *θ*(±Δ*V*)||Δ*V*||, with *θ* being the Heaviside function, which take into account the potentially destabilizing effect due to the creation of holes or stress in the protein interior. The *α*_*i*_(*A*) functions are sigmoids that depend on the solvent accessibility *A* of the mutated residues:

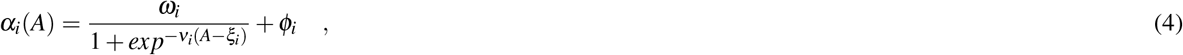

with *ω*_*i*_, *v*_*i*_, ξ_*i*_ and *ϕ*_*i*_ parameters that were identified so as to minimize the mean square deviation between predicted and observed stability changes in a dataset of mutations with experimentally characterized ΔΔ*G*s; note that this dataset has a negligible overlap with **S** (one mutation). For further technical details about the choice of parameters or the construction of the model, see

The classification of the variants of the **S** dataset into deleterious and neutral was performed on the basis of the computed ΔΔ*G* values: SNVs with a predicted ΔΔ*G* higher than a threshold value, which correspond to the most destabilizing mutations, were predicted as deleterious, whereas those with lower ΔΔ*G* were predicted as neutral. The value of the threshold was identified so as to optimize the values of the balanced accuracy (BACC):

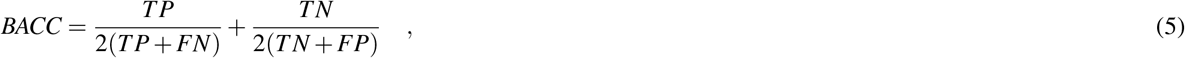

where TP, TN, FP, FN are the true positive, true negative, false positive and false negative predictions, respectively. Positive is chosen to correspond to deleterious variants and negative to neutral ones.

Similarly, we also used the thermal stability predictor HoTMuSiC^35^ for deleterious/neutral classification. This algorithm predicts changes in melting temperature *ΔT*_*m*_ upon mutations rather than changes in folding free energy ΔΔ*G*. These two quantities are anti-correlated only in a first approximation, and yield two different informations about protein stability, as discussed in.^36^ HoTMuSiC uses a distinct combination of the 13 statistical potentials:

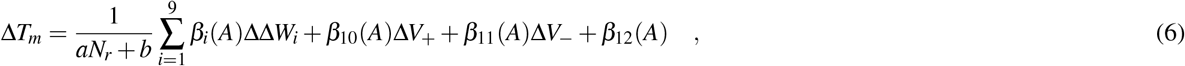

where the *β*_*i*_(*A*) functions are sigmoids of the form specified in Eq. (4), and *N*_*r*_ is the number of residues in the protein. The parameters in the *β*_*i*_(*A*) functions, as well as *a* and *b,* where identified using as cost function the mean square deviation between experimental and predicted Δ*T*_*m*_*s* on a dataset of characterized mutants, that displays only a very small intersection with **S** (3 mutations only). Note that the number of energy terms has been restricted to nine because of the smaller size of the *ΔT*_*m*_ learning dataset, to avoid overfitting.

Using a similar procedure as for the ΔΔ*G* predictions, a given variant of the **S** dataset is considered as deleterious if its *ΔT*_*m*_ value is smaller that a given threshold - which means that the variant provokes a strong thermal destabilization; otherwise it is predicted as neutral. Again, the threshold value was identified using the BACC score as cost function.

#### Stability-based pathogenicity index using an artificial neural network

Using a different combination of the same 13 potentials and two structural features listed in Table S6, we set up a specific predictor of the deleteriousness of a mutation based on stability criteria. It has the peculiarity to relax the hypothesis assumed in the previous section that only destabilizing mutations are likely to be deleterious. Indeed, although most deleterious mutations are destabilizing, some stabilizing mutations can also cause diseases, usually because they affect the protein function, as observed earlier.^37,38^ To take this into account, we considered the stabilizing and destabilizing statistical potential contributions separately, and separated the potential terms into their positive and negative parts:

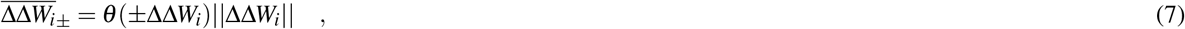

and used them to define the stability-based pathogenicity index I:

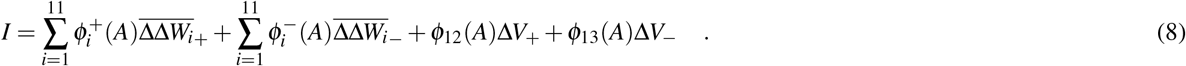

The functions 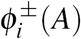 are solvent accessible-dependent sigmoids of the form of Eq. (4). The number of parameters related to the potentials are here almost doubled compared to the PoPMuSiC model structure of Eq. (3).

The model’s parameters were optimized using the double layer artificial neural network (ANN) schematically depicted in Fig. 1a, using as learning set the annotated SNVs from the S dataset and as cost function the mean square deviation between *I* and the annotated variant effect, with deleterious defined as 1 and neutral as 0. The mutations were then considered as deleterious when the pathogenicity index is larger than a given threshold value *ψ* and neutral otherwise. The threshold value was optimized so as to maximize the BACC score (Eq. (5)) of the classification. To evaluate the performance of the method, we used a 5-fold cross-validation procedure. We also tested other validation strategies, among which 10-fold cross validation, with comparable results.

By construction, this new model allows predicting as deleterious both stabilizing and destabilizing mutations. It is a striking illustration that elucidating the biophysical mechanisms underlying the variant effects can be used to guide the model architecture, contribute to improve the classification performances and the understanding of the data.

#### Stability-based pathogenicity index using a probabilistic neural network

Probabilistic neural networks (PNN) introduced in^39^ have architectures that are very different from ANNs, and are utilized for classification purposes. The idea behind their construction consists in a Bayes classification strategy through the estimation of the probability density function (PDF) for each considered category, based on a number of characteristics of the elements to be classified. PNNs are composed of four layers, as shown in Fig. 1b. The input layer contains the features characterizing the considered variant, chosen here to be: 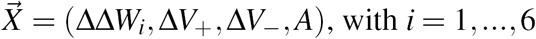 (see Supplementary Material for details). The second layer, called pattern layer, contains as many nodes as there are samples in the training set. Each node, labeled as *j,* contains a probability value computed from the comparison between the feature vectors of the input variant 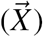 and of the *j*^*th*^ sample 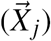, and estimated via a multivariate Gaussian distribution centered on 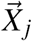:

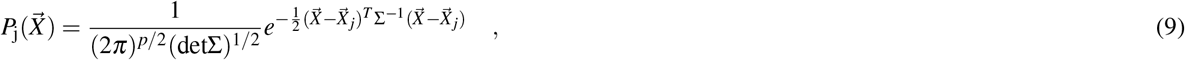

where *p* is the number of features in the input layer and Σ the *p* × p-dimensional covariance matrix of the PDF, considered here to be diagonal. Usually, PNNs use equal covariances for all features, which we do not assume here, hence allowing every feature to contribute with a different weight to the pathogenicity of a mutation. The elements of this Σ-matrix are parameters to be optimized. In the summation layer, the PDFs of the samples that belong to the same category - here neutral or deleterious -are summed, thus defining the category PDFs *P*_*D*_ and *P*_*N*_:

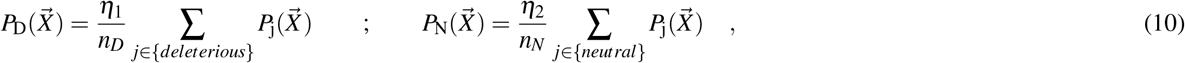

where *n*_*D*_ and *n*_*N*_ are the number of disease-causing and neutral variants in the training dataset, respectively, and *η*_1_ and *η*_2_ two parameters to be optimized. The output layer contains the result of the classification according to the rule that a variant with feature vector 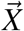 is deleterious if 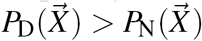 and neutral otherwise. The parameters of the model, *i.e. η*_1_, *η*_2_ and the *p* diagonal elements of the covariance matrix Σ, are identified so as to minimize the BACC score on the variants of the S training set. Again, all performances are evaluated in 5-fold cross validation.

**Figure 1.**
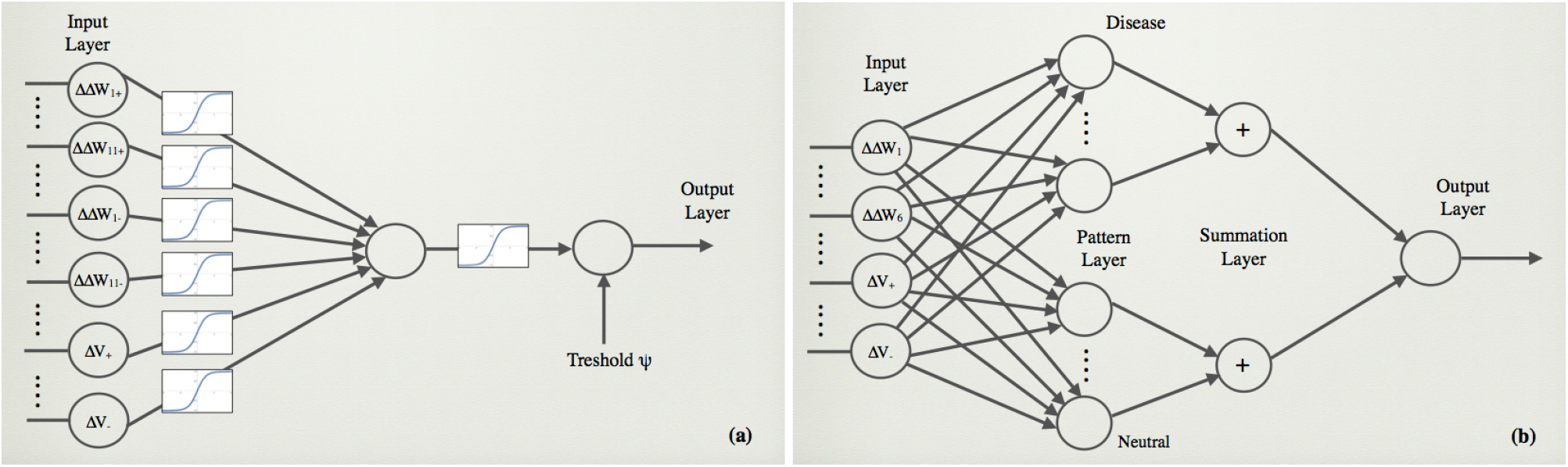
Schematic representation of: (a) the artificial neural network and (b) the probabilistic neural network used in the classification of the variants.

#### Combining stability-based pathogenicity index with evolutionary information

Structural stability and evolutionary information are entangled but distinct notions. On the one hand, biophysical constraints, including stability but also solubility, aggregation propensity, interactions and function, act on natural selection, thus affecting the evolution of proteins, while on the other hand, natural evolution guides mutations, which in turn have an impact on the biophysical properties of proteins.

In our last predictor, both types of information are joined to obtain a more complete picture of the variant effects. We would like to stress that, in contrast to the black-box machine learning approach usually employed in variant classification, we tested concretely whether the stability change due to a residue substitution impacts directly on the variant’s deleteriousness, taking also into account the role of the evolutionary pressure.

The model structure employed is a simple combination of the Provean^34^ score, noted PRO, with the pathogenicity index I defined in Eq. (8). The new index, called 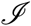, is defined as:

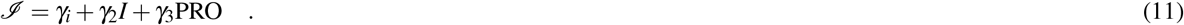

The three coefficients *γ*_*i*_ as well as the classification threshold were identified by optimizing the BACC score on the S dataset. Again, all tests were performed in strict 5-fold cross-validation. We call this predictor SNPMuSiC.

## Results

### Analysis of the statistical potentials and other structural features

Let us start analyzing the solvent accessibility *A* of the mutated residues, which is one of the structural attributes known to be related to variant deleteriousness.^19^ Indeed, mutations in the core are usually more stabilizing or more destabilizing with respect to surface mutations, since buried residues play a special role both in the early folding stages and in the stability of the folded structure. As a consequence, core residue substitutions are more likely to cause structural rearrangements and/or the loss of protein function, and have thus an higher probability to be deleterious for the organism. This is what we observe in Fig. 2a: variants have a higher probability to be disease-causing if the mutation is in a buried region, defined as *A* <20%, while the neutral variant distribution shows only a mild dependence on A with a weak decrease of the probability for large A-values.

**Figure 2.**
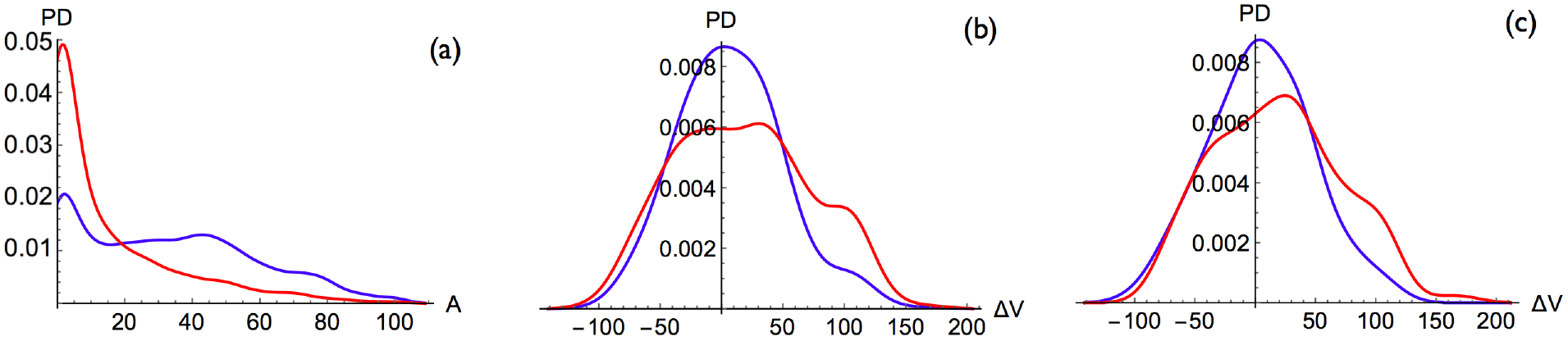
Probability density distributions of deleterious mutations (red curve) and neutral mutations (blue curve) for: (a) the solvent accessibility *A* (0%-100%) of the mutated residues; (b) the change in volume ∆*V* (in *Å*^2^) for residues with *A* ≤60%; (c) the change in volume ∆*V* for residues with *A* >60%; in our conventions, mutations of smaller into larger residues have a positive ∆*V* value.

The second structural feature we considered is the change in volume ∆*V* upon residue substitution. We found that in the totally and partially buried regions, up to a solvent accessibility *A* ≤ 60%, the substitutions from smaller into larger amino acids (positive ∆*V*s) have a higher probability to be deleterious, and so are, albeit to a much smaller extent, the substitutions from larger into smaller amino acids (negative ∆*V*s), as shown in Fig. 2b. In other words, substitutions that create stress or holes in the protein interior are more likely to be deleterious; holes are, however, easier to manage through limited structural rearrangements than stress. On the protein surface, defined as *A* >60%, mutations from smaller to larger residues still have a higher probability to be disease-causing, but less than in the core (Fig. 2c), whereas no difference is observed for substitutions from larger to smaller residues. Note that the differences between deleterious and neutral variant distributions, both for the solvent accessibility and the volume change, are statistically significant, as measured by a very low KS-test P-value (see Methods) shown in Table S6.

Next we compared the probability density functions of deleterious and neutral mutations for the changes in folding free energy ∆∆*W* computed by each of the 13 statistical potentials listed in Table S6. We can classify the results into three classes, whose typical behavior is depicted in Figs 3a-c. In the first class, disease-causing variants are associated with both highly stabilizing and highly destabilizing values. In class 2, the deleterious variants are more frequently associated with destabilizing mutations but not with stabilizing mutations. In the last class, the difference between deleterious and neutral distributions is not statistically significant, as shown by a Kolmogorov-Smirnov (KS) test P-value larger than 0.05. Note that the biophysical interpretation of the neutral and deleterious variant distributions is here less obvious than for solvent accessibility and volume changes, since statistical potentials describe complex interactions between sequence and structure factors.

**Figure 3.**
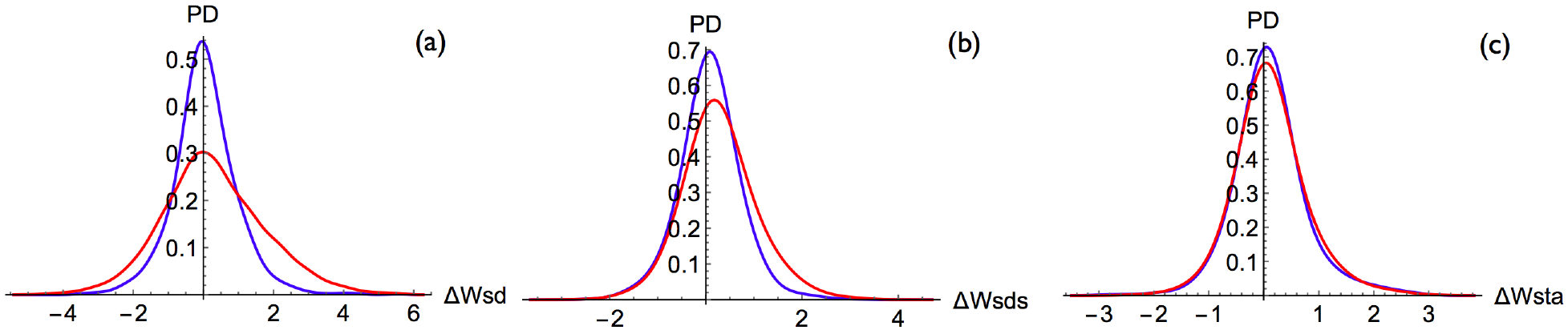
Probability density distributions of deleterious mutations (red curve) and neutral mutations (blue curve) for the changes in folding free energy ∆∆*W* (in kcal/mol) computed with the following statistical potentials: (a) the distance potential ∆*W*_*sd*_; (b) the distance potential ∆*W*_*sds*_; (c) the torsion angle and solvent accessibility potential ∆*W*_*sta*_; in our conventions, positive ∆∆*W* values correspond to destabilizing mutations.

### Classification Performances in Cross Validation

The structural and stability features analyzed in the previous section were used, alone or in combination, to classify single-site mutations into deleterious or neutral, as described in the Methods section. The performance of this classification was evaluated in 5-fold cross validation at the mutation level by training the model on 4/5 randomly chosen mutations that belong to the S dataset and applying it on the remaining entries. This procedure was repeated five times, considering each of the five subsets in turn as test set. The average scores are reported in Table 1. Another 5-fold cross-validation was performed at the protein level, by dividing the proteins rather than the mutations into five subsets; the results are shown in Table S7. No significant differences are observed between the two types of cross-validation scores.

**Table 1.** Performance of the different prediction methods in mutation-based 5-fold cross validation on the learning set. Sensitivity is defined as *TP*/(*TP+FN*), specificity as *TN*/(*TN* + *FP*), positive predictive value (PPV) as *TP*/(*TP* + *FP*), and negative predictive value (NPV) as *TN*/(*TN* + *FN*). Note that the scores and threshold values correspond to averages on the 5-fold cross-validation experiments.

| Method | Sensitivity | Specificity | PPV | NPV | BACC | Threshold | AUROC |
| --- | --- | --- | --- | --- | --- | --- | --- |
| PoPMuSiC | 0.650 | 0.618 | 0.839 | 0.365 | 0.633 | 0.75 kcal/mol | 0.680 |
| HoTMuSiC | 0.593 | 0.648 | 0.838 | 0.341 | 0.620 | -1.8 °C | 0.658 |
| Solvent Accessibility | 0.707 | 0.655 | 0.863 | 0.421 | 0.681 | 18.0 % | 0.719 |
| PNN | 0.692 | 0.717 | 0.883 | 0.431 | 0.705 | - | 0.763 |
| ANN | 0.698 | **0.724** | 0.886 | 0.438 | 0.711 | 0.74 | 0.768 |
| Provean | **0.852** | 0.583 | 0.862 | **0.562** | 0.718 | -2.5 | 0.800 |
| SNPMuSiC | 0.789 | 0.712 | **0.894** | 0.524 | **0.751** | 0.66 | **0.825** |

Strikingly, the variant classification based solely on solvent accessibility gives quite good results, with a BACC score of 0.68, as expected by the large dissimilarity between deleterious and neutral variant distributions shown in Fig. 2a. This result again demonstrates the important role of core residues in folding, structure and stability, and their sensitivity to mutations.

The classifications based on the prediction of thermodynamic and thermal stability changes upon mutations computed by PoPMuSiC (Eq. (3)) and HoTMuSiC (Eq. 6)), respectively, yield BACC scores of 0.62 and 0.63; the neutral and deleterious variant distributions for PoPMuSiC are shown in Fig. 4a. There is thus a significant, but limited, correlation between destabilization and deleteriousness. Two reasons can be invoked to explain why this correlation is not higher. The first is that deleteriousness can be caused by destabilization but also by other factors. The second explanation is that not only destabilizing but also stabilizing mutations can be deleterious, as seen in Fig. 3a. To illustrate this important point, we describe some disease-causing variants associated with a gain in structural stability in the next subsection.

**Figure 4.**
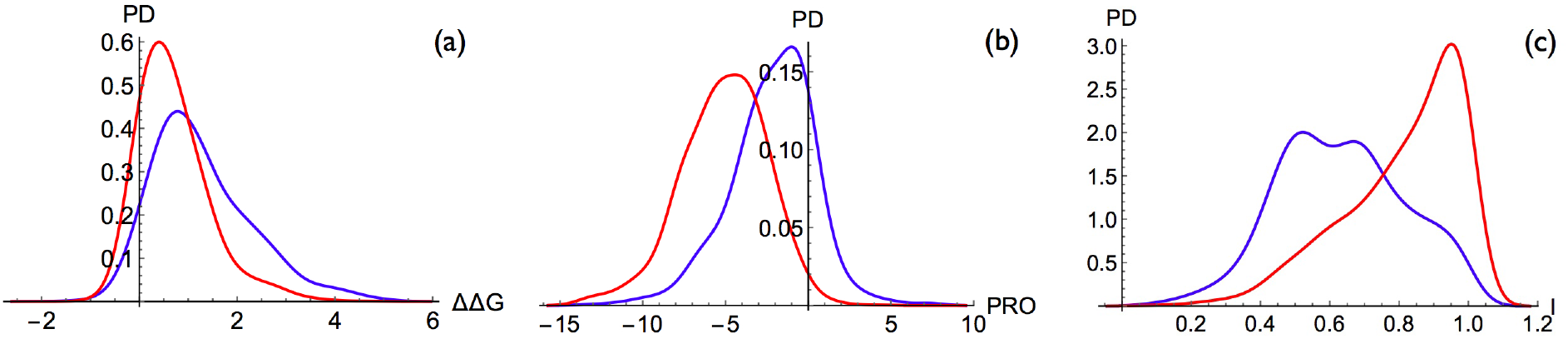
Probability density distributions of deleterious mutations (red curve) and neutral mutations (blue curve) for (a) the change in folding free energy ∆∆*G* computed by PoPMuSiC (Eq. (3)) (in kcal/mol), (b) the Provean score,^3,4^ and (c) the pathogenicity index *I* computed by the ANN model (Eq. (8)).

To take into account that both stabilizing and destabilizing mutations can cause diseases, we set up two new classifiers that use the same structural features and statistical potentials as PoPMuSiC and HoTMuSiC but are based on two different neural network architectures, an ANN (Eq. (8)) and a PNN (Eq. (10)). These two classifiers both reach a BACC score of 0.71; the neutral and deleterious variant distributions of the ANN model are shown in Fig. 4c. As expected, these two models take more properly into account the weight of the different energy contributions in the determination of the variant deleteriousness. The fact that the ANN and PNN model structures yield the same score suggests that it corresponds to the maximum performance that can be obtained from the features considered.

Quite interestingly, the Provean classifier^3, 4^ reaches the same score as our ANN and PNN models (0.07 higher), though the former is based on evolutionary information that mixes stability with other biophysical characteristics such as solubility and activity while the latter purely exploits stability information. The variant distributions obtained with this predictor are depicted in Fig. 4c. A closer analysis shows, however, some important differences. Our ANN and PNN prediction methods have a sensitivity of about 0.70 that is much lower than the Provean value of 0.85, and a specificity of 0.72 that is much higher than the Provean value of 0.58 (Table 1). The high specificity and low sensitivity of our ANN and PNN classifiers come from their number of false positives being reduced by nearly 50% compared to Provean, as well as a larger false negative rate. This result can be explained as follows: variants predicted as strongly stabilizing or destabilizing are usually well classified as disease-causing as they are very likely to have an impact on the phenotype; this accounts for the low false positive rate. Conversely, our ANN and PNN predictors are unable, by construction, to predict the deleterious effects due to biophysical characteristics other than stability; this rationalizes the high number of false negatives.

Our last and best predictor, that we call SNPMuSIC, combines the deleteriousness index *I* of the ANN model (Eq. (8)) with the Provean score, thus defining the 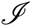 index (Eq. (11)). This model reaches the highest BACC score, *i.e.* 0.75, and the highest Area Under the Receiver Operating Characteristic curve (AUROC score), *i.e.* 0.83; the associated distributions of neutral and deleterious variants are quite well separated as shown in Fig. 5. This predictor has BACC and AUROC scores that are more than 3% higher than both Provean and ANN, which is statistically significant (P-value < 10^−4^ using a bootstrap test). It performs even better than more complete classification methods, based on a whole range of sequence and structural features, such as Polyphen-2.^10^ Strikingly, SNPMuSiC increases the sensitivity of the ANN and PNN models up to 0.79, and almost maintains their high specificity value (0.71) (Table 1).

**Figure 5.**
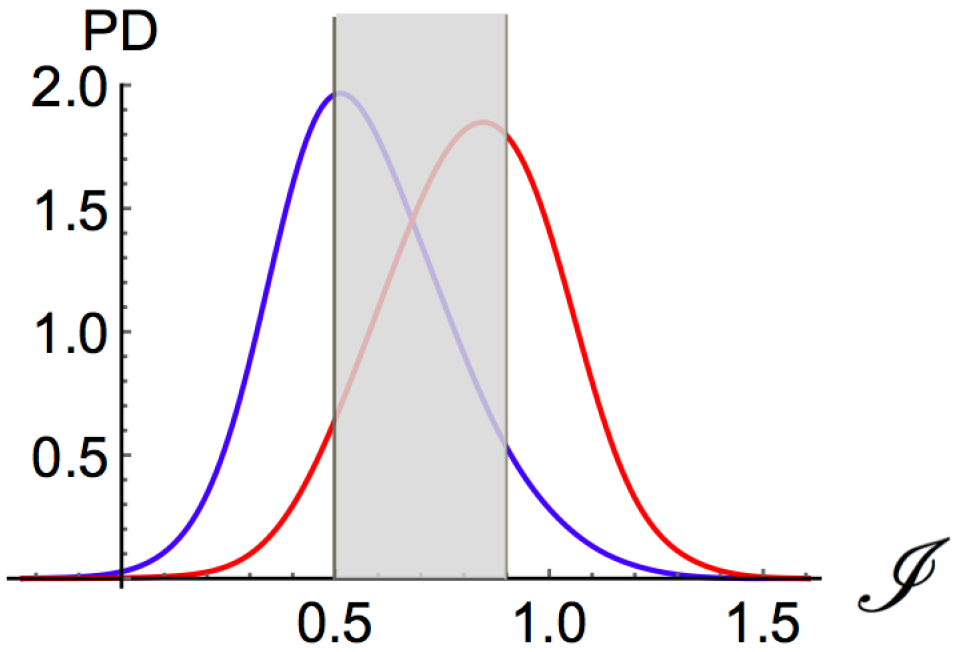
Probability density distribution of deleterious mutations (red curve) and neutral mutations (blue curve) for the pathological index 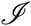 (Eq. (11)) computed by SNPMuSiC. The distribution curves in the high confidence intervals, which lie from either side of the two vertical lines, are depicted on a white background.

SNPMuSiC will be available soon on the PoPMuSiC and HoTMuSiC webserver at https://soft.dezyme.com/.

### Classification Performances on the Test Sets

To analyze in more detail the performances of our predictors, we applied them to four different test sets **S**_**H**_, **S**_**NH**_, **S**_**CV**_ and **S**_**SSC**_ taken from^12^ and specified in Tables S2-S5. In contrast to our learning set, these sets do not contain well resolved X-ray structures but instead structures obtained by comparative modeling. This test amounts thus to evaluating the performance of our predictors on low-resolution structures. They were compared to commonly used deleteriousness predictors that do not include contextual information, *i.e*. Polyphen-2,^10^ Provean,^3,4^ CADD,^9^ Sift^11^ and VIPUR.^12^ The results of these programs are taken from^12^ when available.

The first two test sets correspond to human variants from HumVar^10^ and to non-human variants, respecively. We excluded from the former set the variants present in the training sets of our method and of Polyphen-2, as well as the proteins whose 3D structure is modeled from templates that have less than 30% identity with respect to the target sequence, in order to avoid incorrectly modeled structures. The results are shown in Table 2.

**Table 2.** Comparison of the performances of the different predictors on the test sets **S**_**H**_ and **S**_**NH**_. These datasets have no overlap with the training sets of Polyphen-2, the solvent accessibility-based method, PoPMuSiC and SNPMuSiC. For VIPUR, the scores for **S**_**H**_ are those labelled as being cross validated in,^12^ while for the **S**_**NH**_ set, no cross validated scores are available. For the sequence-based methods Provean, the dataset overlap has not been considered, although it plays a role in the identification of the threshold values. See Table 1 for further details.

| Method | Sensibility | Specificity | PPV | NPV | BACC |
| --- | --- | --- | --- | --- | --- |
| **Human Variants** |  |  |  |  |  |
| PoPMuSiC | 0.539 | 0.621 | 0.785 | 0.344 | 0.580 |
| Solvent Accessibility | 0.584 | **0.655** | 0.813 | 0.380 | 0.620 |
| VIPUR | 0.868 | 0.553 | **0.833** | 0.620 | **0.710** |
| Polyphen-2 | **0.948** | 0.330 | 0.784 | 0.712 | 0.639 |
| Provean | 0.943 | 0.387 | 0.798 | **0.727** | 0.665 |
| SNPMuSiC | 0.759 | 0.603 | **0.831** | 0.494 | 0.681 |
| **Non-Human Variants** |  |  |  |  |  |
| PoPMuSiC | 0.531 | 0.543 | 0.734 | 0.328 | 0.537 |
| Solvent Accessibility | 0.553 | **0.636** | 0.782 | 0.375 | 0.594 |
| Polyphen-2 | **0.964** | 0.304 | 0.771 | **0.778** | 0.634 |
| Provean | 0.940 | 0.339 | 0.771 | 0.703 | 0.640 |
| SNPMuSiC | 0.782 | 0.539 | **0.801** | 0.511 | **0.661** |

On both test sets, the solvent accessibility-based predictor has the highest specificity and Polyphen-2 the largest sensitivity. Hence, the former has the largest proportion of neutral mutations that are correctly identified as such, while the second has the largest proportion of correctly identified deleterious mutations. The SNPMuSiC predictor has the largest BACC and PPV scores for the non-human test set. On the human set, VIPUR has a larger BACC with respect to the other tools while VIPUR and SNPMuSiC have a similar PPV.

We also evaluated the performances on the two other sets **S**_**CV**_ and **S**_**SSC**_, which contain neutral and disease-causing human variants from ClinVar^26^ and *de novo* missense mutations of the Simon Simplex Collection (SSC),^27^ respectively. The results are shown in Table S8. Here, we could not filter out incorrect 3D models, as we do not have access to the sequence identity between the target and template sequences. This could explain why sequence-based predictors sometimes perform better than structure-based ones. In spite of this, the PPV score of SNPMuSiC is the highest on both sets. Moreover, the BACC score is the highest on the **S**_**CV**_ set compared to all other predictors, and on the **S**_**SSC**_ set compared to the other structure based methods Polyphen-2 and Vipur.

This leads us to the conclusion that SNPMuSiC is also applicable to mutations in proteins with low-resolution (modeled) structures, which drastically increases its application possibilities. Moreover, in comparison with a series of other widely known classification tools, it has the highest BACC for several test sets and the highest PPV for all sets analyzed. Hence, the deleterious mutations predicted by SNPMuSiC have a large chance of being so. This is in accordance with its construction: it only predicts mutations that are deleterious because of stability reasons, and misses some other deleterious mutations that instead are caused by other biophysical mechanisms.

### Link between protein stabilization and disease

While destabilizing mutations can clearly be deleterious as they are likely to cause structural modification which can negatively impact on protein function, it is less obvious that variants with a stabilizing effect can also be disease causing. To clarify this point we analyzed the ten deleterious mutations of the variant dataset **S** that are predicted to be the most thermodynamically stabilizing by PoPMuSiC; they are reported in Table 3. All the mutated residues are located in the protein core, with an accessibility lower than 15%. Note that PoPMuSiC’s ∆∆*G* and HoTMuSiC’s ∆*T*_*m*_ prediction values are not always anti-correlated, an illustration of the imperfect relation of thermodynamic and thermal stabilities.^36^

**Table 3.** List of the 10 mutations from the S dataset which are annotated as disease causing and are predicted as the most stabilizing (*i.e.* with the most negative ∆∆*G* values) by PoPMuSiC. Light blue cells mean variants predicted as neutral by the classifier mentioned in the column’s title, while red cells contain variants classified as deleterious.

| Protein | Chain | Mutation | Biophysical effect | $\Delta\Delta G$<br>PoPMuSiC<br>(kcal/mol) | $\Delta T_m$<br>HoTMuSiC<br>(°C) | $I$ index<br>ANN | $\mathcal{J}$ index<br>SNPMuSiC |
| --- | --- | --- | --- | --- | --- | --- | --- |
| 1m6i | A | E493V | Increase of NADH affinity | -2.24 | 1.7 | 1.00 | 4.8 |
| 2wzb | A | D163V | Decrease of enzymatic activity | -1.62 | 1.0 | 1.00 | 4.1 |
| 3hcn | A | T283I | Decrease of enzymatic activity | -1.46 | 1.1 | 0.92 | 1.9 |
| 2izz | A | G206W | Loss of function | -1.35 | 2.8 | 1.00 | 1.4 |
| 2nt0 | A | D399Y | Decrease of enzymatic activity | -1.31 | 0.8 | 1.00 | 4.4 |
| 3f9m | A | G385V | Decrease of enzymatic activity | -1.23 | -0.2 | 0.84 | 1.3 |
| 4do4 | A | R329W | Decrease of enzymatic activity | -0.99 | 0.3 | 0.99 | 2.3 |
| 1aly | A | G227V | Loss of ligand binding | -0.96 | 1.8 | 0.99 | 1.7 |
| 4az3 | A | S23Y | Loss of phosphorylation | -0.95 | 1.8 | 1.00 | 0.7 |
| 2nt0 | A | D380H | Decrease of enzymatic activity | -0.86 | 0.3 | 1.00 | 3.1 |

Obviously, the PoPMuSiC and HoTMuSiC-based classifiers wrongly predict these ten mutations as neutral, since only destabilizing mutations have a chance to be classified as deleterious by these two models. In contrast, both our ANN-based predictor and SNPMuSiC, which are designed to be able to predict stabilizing disease-causing mutations, correctly classify all ten mutations as deleterious.

It is instructive to briefly analyze the biophysical mechanisms that are involved in the pathogenic phenotype of these SNVs caused by protein stabilization. These are taken from the annotations of the variants reported in the Uniprot database.^23^ As shown in Table 3, the protein stabilization leads to a decrease of the enzymatic activity in the majority of the cases, which can be explained by the mutation being close to the protein active site or inducing inhibiting allosteric effects. Other mutations cause the change in affinity for ligands or protein partners^41,43^ or affect post-translational modifications sites.^44^

In summary, this investigation shows that biophysical knowledge at the molecular level - in this case, the fact that both strongly stabilizing and destabilizing mutations are likely to cause diseases - can guide the design of the model structures, improve the classification performances and lead to a deeper understanding of the variant effects.

### Reliability of structural stability prediction

Our SNPMuSiC method reaches the very high positive predicted value (PPV) of 89%, as shown in Table 1, which indicates that when a mutation is predicted as deleterious, it has a large probability of being so. The negative predictive value is much lower, 52%, indicating that when a mutation is predicted as neutral, it has still 50% chance of being disease-causing, albeit for a reason that is not related to stability.

The idea here is thus to identify a subset of mutations, whose pathogenicity index 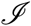 falls above a certain confidence threshold and whose prediction reaches an even higher level of accuracy. For these mutations, the protein stability is the main contributing factor to the variant’s deleteriousness. We also identified another subset of mutations, with 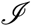 below another confidence threshold, which are predicted as neutral with a good accuracy. This strategy is quite useful in the perspective of combining SNPMuSiC with other variant effect predictors that are based on different biophysical characteristics such as aggregation propensities or flexibility.

Setting the 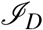 confidence threshold equal to 0.9, we find that the PPV reaches 95% on the subset **S**_*high*_ with 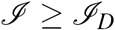. This subset contains about 1,500 mutations, which corresponds to 27% of the whole S set. Here we can be almost sure of the deleteriousness of the mutations and of its molecular cause. Indeed, combining the SNPMuSiC prediction with the outputs of PoPMuSiC and HoTMuSiC, we can determine if they are deleterious due to thermodynamic or thermal stabilization or destabilization.

With the 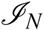 threshold for neutral mutations equal to 0.5, we obtain an NPV of 72% on the subset **S**_*low*_ for which 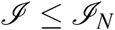. This subset contains about 700 mutations, thus about 13% of the total **S** set. For these mutations, we have a good indication of their neutral impact on the phenotype.

The BACC score computed on the subsets characterized by 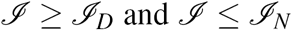, representing 40% of the **S** set, is also significantly improved and reaches 0.87. The SNPMuSiC distribution for deleterious and neutral variants at both sides of the confidence thresholds are shown in Fig. 4.

## Conclusion

Here we made a further step towards the prediction of the biophysical causes of the deleteriousness of non-synonymous SNVs. We focused on protein stability which guides a series of crucial biophysical mechanisms that encompass folding, interactions, and function. Our analysis can be extended and generalized to all other types of biophysical characteristics that play a role at the protein level and could be at the basis of deleteriousness, such as solubility, aggregation, allostery, flexibility, and catalytic activity. This will lead to a complete view of the relation between the effect of SNVs at the molecular level and disease, and pave the way towards personalized medecine.

Let us give a flavor of the next stages of our quest. We will include the changes caused by mutations on the protein aggregation properties, which are not directly related to protein stability^40^ even though they are frequently considered as correlated. Other factors that we will add to this analysis is the SNV effect on protein activity and on protein-protein interaction networks,^41,42^ which are undoubtedly important factors in the initiation of diseases.

Finally we would like to underline the importance of using the 3D structures of the proteins in which SNV occur to predict and interpret their biophysical and disease-causing effects. This data is essentially overlooked in most commonly used deleteriousness classification methods, which drastically limits their interpretative power. Two reasons can be invoked to explain this shortcoming. On the one hand, protein sequences are much easier to handle than protein structures, and on the other hand, many proteins have no available experimental structure. This last issue has, however, to be relativized, as many protein structures can be obtained through comparative modeling. Given that our methods use coarse-grained representations of protein structure, they can be applied on protein models with only a small loss of accuracy.^45^ We would like to end by stressing the broad applicability of structure-based predictors, as it has been estimated that almost half of all structured proteins have either an experimental or reliably modeled structure.^46^

## Supplementary Information

- Non-synonymous SNV training set (Table S1)
- Non-synonymous SNV test sets (Table S2-S5)
- Statistical potentials (Table S6)
- Performance of our predictors in cross validation on the training set (Table S7)
- Performance comparisons on the test sets (Table S8)

## Acknowledgements

We thank Georges Coppin, Suchsia Chao and Thanh Phuong Pham for participating in the early stages of this project, and Jean Marc Kwasigroch, Daniele Raimondi and Andrea Gazzo for help in setting up the training dataset.

## Author contributions statement

F.P. and M.R. conceived the experiments, F.A., F.P. and M.G. conducted the experiments, F.A., F.P. and M.R. analyzed the results, F.P. and M.R. wrote the paper. All authors reviewed the manuscript.

## Additional information

The authors declare no competing financial interests.

